# Healthcare professionals and students’ awareness of Chagas disease: design and validation of ChaLKS (Chagas Level of Knowledge Scale)

**DOI:** 10.1101/741488

**Authors:** José M. Ramos-Rincón, José J. Mira-Solves, Violeta Ramos-Sesma, Diego Torrús-Tendero, Jara Llenas-García, Miriam Navarro

**Affiliations:** Department of Clinical Medicine, Universidad Miguel Hernández de Elche, Campus de Sant Joan d’Alacant, Alicante, Spain; Department of Internal Medicine, Hospital General Universitario de Alicante, Instituto de Investigación Sanitaria y Biomédica de Alicante (ISABIAL-Fundación FISABIO), Alicante, Spain; Department of Health Psychology, Universidad Miguel Hernández de Elche, Campus de Elche, Alicante, Spain; Health Psychology Unit, Departamento de Salud Alicante-San Juan, Alicante, Spain; Department of Internal Medicine, Hospital Universitario de Torrevieja, Alicante, Spain; Parasitology Area, Universidad Miguel Hernández de Elche, Campus de Sant Joan d’Alacant, Alicante, Spain; Department of Internal Medicine, Hospital Vega Baja, Fundación para el Fomento de la Investigación Sanitaria y Biomédica de la Comunitat Valenciana (FISABIO), Orihuela, Alicante, Spain; Department of Public Health, Science History and Gynecology. Universidad Miguel Hernández de Elche, Campus de Sant Joan d’Alacant, Alicante, Spain

**Keywords:** Chagas disease, awareness, knowledge scale, students, physicians, pharmacists, Spain, Europe, Latin America

## Abstract

**Background:** There are few studies evaluating awareness of Chagas disease among healthcare professionals attending migrants from Latin America or working in Chagas-endemic areas. The objective of this study was to design and validate instruments for assessing knowledge about Chagas disease among healthcare students and residents as well as students and professionals of social and other health science disciplines.

**Principal findings:** Two validated scales were obtained: the 10-item Chagas Level of Knowledge Scale for healthcare professionals (ChaLKS-Medical) and the 8-item ChaLKS-Social&Health for potential aid workers from those fields. Both scales were considered adequate in terms of readability, internal consistency, construct validity and discriminant validity.

The mean number of correct answers on the ChaLKS-Medical among respondents from non-healthcare versus healthcare sectors was 1.80 (standard deviation [SD] 2.55) versus 7.00 (SD 2.32) (p<0.001). The scores on the ChaLKS-Social&Health also discriminated between the knowledge levels in these two groups (1.76 [SD 2.47] versus 6.78 [SD 1.55], p<0.001). Knowledge among medical/pharmacy students and residents on the ChaLKS-Medical was acceptable (mean 5.8 [SD 2.1] and 7.4 [SD 2.2], respectively; p<0.001). Respondents’ knowledge on Chagas disease was greater in those who had previously received information on the disease; this was true in both respondents from the healthcare sector (mean correct answers, ChaLKS-Medical: 7.2 [SD 2.1] versus 4.3 [2.6], p<0.001) and in potential aid workers (mean correct answers, ChaLKS-Social&Health: 5.1 [SD 2.5] versus 1.1 [SD 1.9], p=0.001).

**Conclusions:** The metric properties of both scales are adequate for their use in supporting aid operations in Chagas-endemic countries or in providing health and social care to migrant populations in non-endemic countries. The results of these scales also provide orientation regarding the knowledge gaps to be filled in future health professional’s training programs.

**Author summary:** Chagas disease is endemic to much of Latin America, but it is still largely unrecognized in Europe, in part because of the lack of professional training in tropical medicine and global health among healthcare professionals. Few studies have evaluated awareness of Chagas disease among healthcare professionals and medical students, and to our knowledge none have done so in aid workers or volunteers working with Latin American populations. The aim of our study was to design and validate instruments for assessing knowledge about Chagas disease among healthcare students and professionals as well as potential aid workers. Two validated scales were obtained: the Chagas Level of Knowledge Scale for healthcare professionals (ChaLKS-Medical) and the ChaLKS-Social&Health for potential aid workers in social and other health fields. The two scales could be especially useful to reinforce training on Chagas disease among medical students and health and social aid workers working in Chagas-endemic regions or with migrants from those countries, regardless of whether their mission is related to Chagas or not. Moreover, this could reinforce the cross-cutting nature of programs in areas endemic to Chagas and other neglected tropical diseases.

## INTRODUCTION

Chagas disease (CD) is a parasitic (*Trypanosoma cruzi*), mainly vector-borne disease with a high public health impact in Europe, where other transmission routes (blood transfusion, transplantation, mother-to-child) may also occur and are not completely controlled [1]. Rising immigration from Latin America to Europe and especially to Spain has led to an increase in cases, mainly in asymptomatic immigrants in the chronic phase of the disease [2]. However, CD is still largely unrecognized in Europe, and fewer than 10% of cases are diagnosed [3]. This has led to active efforts to identify patients in order to bring them into the healthcare system [4].

Under-diagnosis of CD in Europe has three main component causes, related to the population at risk (lack of knowledge and awareness about the disease, fear, stigma, barriers to access healthcare system); healthcare professionals (lack of training in tropical medicine, global health and cultural diversity in the consultation); and public health measures (so far, insufficient to address the challenge of detecting and controlling this emerging neglected tropical disease [NTD]) [1,5].

Surveys in the United States show a general lack of awareness of CD among physicians across specialties [6-8]. Studies have indicated that US physicians may not consider CD when diagnosing immigrant patients from Chagas-endemic areas [8]. The American Academy of Cardiology has recognized the need to train its physicians in CD in order to improve care in a context where a growing proportion of the patient population is affected [9]. In Europe, there are few studies evaluating awareness of CD among healthcare professionals [10,11], and little is known about medical students’ knowledge on the issue [12,13].

At the same time, international aid work in Latin America has changed drastically in the past decade, also affecting social organizations. This change is also relevant from the Spanish perspective [14,15]. However, similarly to the health sector, there are scarce data about aid workers’ and volunteers’ knowledge on CD prior to working with populations in or from endemic regions.

The objective of this study was to design and validate instruments for assessing knowledge about CD among medical and pharmacy students and residents, and among students of other health science and social disciplines who are interested in international cooperation in Latin America or in attending Latin American migrants.

## MATERIAL AND METHODS

### Study design

Cross-sectional study conducted in Alicante, Spain, between January 2016 and March 2018, focused on the validation of both scales.

### Participants

We recruited a convenience sample of 349 participants (66 students in fields other than medicine or pharmacy who were interested in aid work, plus 283 medicine and pharmacy students and residents). The sample size was determined to detect a difference of 0.39 points, with a confidence level of 95%. Inclusion criteria were: medicine and pharmacy students; students and professionals from other disciplines related to aid work who had obtained their bachelor’s degree in Europe; and doctors, pharmacists, and other health professionals who had recently completed or were completing their specialty training in Spain. Participants were recruited through six events or channels (conferences or classes on aid work, etc.: **Supplementary table S1)** and replied using a self-administered questionnaire (pencil-paper and online approaches were used). All participants were informed about the objective of the study before completing the survey.

To perform the study, a reactive questionnaire, based on diverse sources of information, was designed to assess respondents’ level of knowledge. The tool was analyzed for readability, reliability, and validity.

### Scale design

Two authors developed an initial set of questions based on a questionnaire aimed at populations at risk of CD [16]. It had 26 reactive items, encompassing knowledge on Chagas epidemiology (5 items), transmission routes (8 items), clinical characteristics (6 items), diagnosis (4 items), and treatment (3 items) (**see supplementary table S2**). This proposal was assessed for relevance, clarity, and priority by three internists, three family and community medicine specialists (all with experience in CD), and three physicians from Latin America with knowledge of CD, in order to best discriminate between different levels of knowledge (face and content validity analyses). These experts ranked the importance of the items and recommended eliminating three. Following this analysis, our team agreed to design a test on CD directed toward medical and pharmacy students and another directed toward students in other health and social fields who were interested in international aid work in areas where the population was at risk of CD.

The version for healthcare students and residents included 23 items, while the one for potential aid workers had 15 items in Spanish after eliminating 8 items on diagnosing and treating Chagas (**Supplementary material: table S3a in Spanish and table S3b in English**). All questions for both questionnaires included three possible responses: “yes,” “no,” and “I don’t know.” The scores in both questionnaires were the sum of the correct answers; incorrect answers were not penalized. One of the questions on transmission was initially formulated as an open question (*What insect spreads Chagas disease?*). The four most frequent answers were: mosquito, sandfly, fly, and kissing bug (S**upplementary material: table S2**). In the final version of the questionnaire, the item was reformulated as closed in order to maintain the same format as for the other items (*Do kissing bugs spread Chagas disease?*) (**Supplementary material: tables S5a and S5b**).

To assess comprehension and the potential for misunderstandings, we asked three third-year students in occupational therapy, podiatry and physical therapy, plus three last-year medical students, to read the respective versions of the questionnaire for aid workers and health professionals. These student profiles were chosen because they are used to answering multiple-choice questions and would thus be able to detect poorly formulated questions more easily.

The difficulty index of each question was analyzed by dividing the number of correct responses by the number of total respondents; higher scores corresponded to easier items. We deemed indexes of 0.50 to 0.60 to be an average level of difficulty; indexes of 0.40 to 0.70 to be a moderate level of difficulty; we also aimed to include 5% of questions with a high level of difficulty (less than 0.30).

### Reliability

Cronbach’s alpha was applied as measure of internal consistency. Reliability was calculated using the Split-half method, applying the Spearman-Brown coefficient. A minimum of 0.70 was considered acceptable for both statistics. The item-total correlation was also analyzed to characterize the metric properties of the elements, excluding those elements with low correlations. A minimum coefficient of 0.35 for Pearson’s correlation was considered acceptable.

### Construct validity

An exploratory factor analysis (EFA) was performed to determine the factorial structure of each instrument using the Principal Components technique, followed by the Varimax rotation. A factor loading greater than 0.5 was considered an acceptable level of missing data. Prior to this analysis, the Kaiser-Meyer-Olkin Measure of Sampling Adequacy was applied to identify the proportion of variance in the items that might be caused by underlying factors. Also, Bartlett’s test of sphericity tests was applied to determine whether the exploratory factor analysis was suitable for structure detection.

### Discriminant validity

The responses from the two sub-samples were compared to check that healthcare students and resident physicians consistently obtained better scores than non-healthcare students (discriminant validity). Using the student’s t test, we compared the number of correct answers obtained by respondents who reported having versus not having received prior training in CD, and between those who had received their obtained their bachelor’s degree in Latin America versus Europe. We also compared the performance of different subgroups: medicine and pharmacy students along with medical and pharmacy residents. Finally, we compared the scores of potential aid workers and students, and medical and pharmacy professionals.

### Translation

An English-native translator with expertise in scientific documents translated the scales from Spanish. A forward-back-translation was then performed by another independent translator to obtain a Spanish version of these instruments (from the English-language ones), in order to check their similarity.

### Ethical aspects

All participants provided informed consent. Those participants who attended the seminars/courses gave their oral consent before the lectures. The study was approved by the Miguel Hernández University’s Project Evaluation Committee (Ref: DMC.JRR.01.16).

## RESULTS

### Scale validation

#### Description of the study sample

**Table 1** presents the characteristics of the study sample, which was made up of 283 students and residents in medicine and pharmacy, plus 66 potential aid workers.

**Table 1.** Characteristics of the two samples of students and professionals in whom the scale was validated

| Characteristics | Medicine and pharmacy students & residents |  | Social and health science* students |  |
| --- | --- | --- | --- | --- |
|  | Mean | SD | Mean | SD |
| Age in years | 28.9 | 7.0 | 21 | 6 |
| Studies | N | % | N | % |
| Medical students | 69 | 19.8 |  |  |
| Pharmacy students | 14 | 4.0 |  |  |
| Pharmacy residents | 18 | 5.2 |  |  |
| Medical residents | 182 | 35.8 |  |  |
| Specialist in family and community medicine | 82 | 9.1 |  |  |
| Specialist in internal medicine and other specialties | 100 | 28.7 |  |  |
| Other students |  |  |  |  |
| Physiotherapy |  |  | 24 | 36.4 |
| Occupational therapy |  |  | 15 | 22.7 |
| Podiatry |  |  | 9 | 13.6 |
| Psychology |  |  | 8 | 12.1 |
| Nursing |  |  | 6 | 9.1 |
| International relations |  |  | 2 | 3.0 |
| Health technology |  |  | 2 | 3.0 |
| **Previous information on Chagas disease** | 255 | 90.1 | 13 | 19.7 |
| **Recruitment events** |  |  |  |  |
| Conferences on cooperation | 66 | 23.3 | 66 | 100 |
| Master in infectious diseases | 47 | 16.6 |  |  |
| Seminar on Chagas disease | 21 | 7.4 |  |  |
| Course on care for immigrant patients | 82 | 29.0 |  |  |
| Pre-congress workshop | 36 | 12.7 |  |  |
| Online | 31 | 11.0 |  |  |
| **Place of birth** |  |  |  |  |
| Spain | 251 | 88.7 | 62 | 93.9 |
| Latin America | 25 | 8.8 | 0 |  |
| Other European country | 6 | 2.1 | 3 | 4.5 |
| Africa | 1 | 0.4 | 1 | 1.5 |

SD: standard deviation.
\*other than Medicine and Pharmacy.

#### ChaLKS for healthcare professionals (ChaLKS-Medical)

Once the elements with saturation in more than one factor or with factorial saturation of less than 0.50 were eliminated, the EFA converged into one dimension with 10 items (**table 2**). The version of ChaLKS that emerged included the items that the experts had considered essential (**Supplementary material: tables S4, S5a, and S5b**). The difficulty indexes ranged from 0.23 to 0.91 (**table 2**). Seven of the 10 items showed an average difficulty index, and 1/10 was deemed moderately difficult. The item-total correlations ranged between 0.30 and 0.53, and Cronbach’s alpha was 0.75. Just over half (54.6%) of the respondents answered seven or fewer items correctly; 16% obtained a perfect score.

**Table 2.** Results of the exploratory factor analysis and difficulty index for the ChaLKS items

| Questionnaire items | ChaLKS-Medical |  | ChaLKS-Social&Health |  |
| --- | --- | --- | --- | --- |
|  | Factorial saturation | Difficulty index | Factorial saturation | Difficulty index |
| Is CD endemic to the Americas? | 0.81 | 0.91 | 0.79 | 0.20 |
| Is CD endemic to Africa? | 0.80 | 0.84 | 0.71 | 0.20 |
| Is CD endemic to Europe? | 0.74 | 0.91 | 0.87 | 0.32 |
| Can CD be transmitted through blood transfusions? | 0.71 | 0.75 | -- | -- |
| Can CD be transmitted from mother to child? | 0.70 | 0.76 | -- | -- |
| Can CD be transmitted by kissing bugs? | 0.68 | 0.59 | 0.66 | 0.15 |
| Can CD affect the heart? | 0.62 | 0.68 | 0.82 | 0.41 |
| Can CD affect the stomach and bowels? | 0.62 | 0.65 | -- | -- |
| Is CD diagnosed via serology? | 0.58 | 0.67 | -- | -- |
| What drug is used to treat CD? | 0.53 | 0.23 | -- | -- |
| Is there treatment for CD? | -- | -- | 0.80 | 0.33 |
| Can CD be transmitted by living with a person who has the disease? | -- | -- | 0.79 | 0.23 |
| Can CD be transmitted through sexual contact? | -- | -- | 0.73 | 0.20 |
|  | Explained variance |  | Explained variance |  |
|  | 46.66% |  | 59.86% |  |
|  | Alpha Cronbach 0.75 |  | Alpha Cronbach 0.89 |  |

CD: Chagas disease

#### ChaLKS for potential aid workers in social and health fields (ChaLKS-Social&Health)

Once the elements with saturation in more than one factor or with factorial saturation of less than 0.50 were eliminated, the EFA converged into one dimension with 8 items. Difficulty indexes ranged from 0.15 to 0.41 (**table 2**). Three of the 8 items (37.5%) showed an average difficulty index and 5/8 (62.5%), a moderate one. The item-total correlations ranged between 0.47 and 0.79. Cronbach’s Alpha was 0.89. The vast majority (87.1%) of the sample answered 5 or fewer questions correctly; 3.2% obtained a perfect score (**table 3**).

**Table 3.**
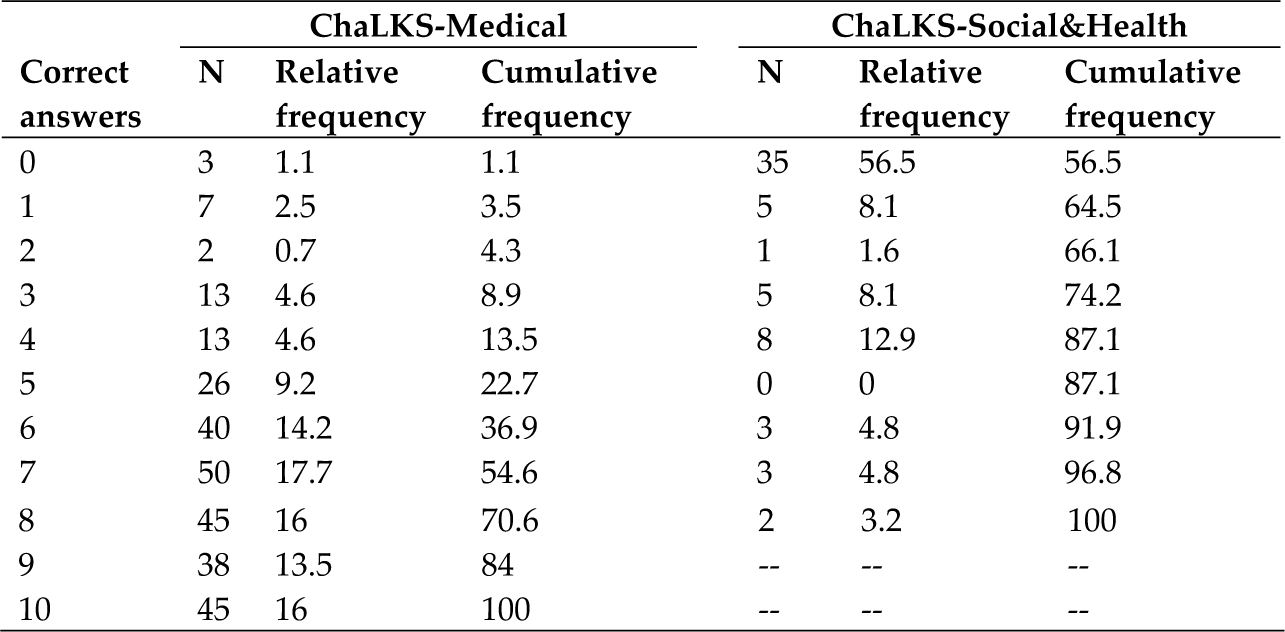
Correct answers on the ChaLKS questionnaire: absolute, relative, and cumulative frequency

| Correct answers | ChaLKS-Medical |  |  | ChaLKS-Social&Health |  |  |
| --- | --- | --- | --- | --- | --- | --- |
|  | N | Relative frequency | Cumulative frequency | N | Relative frequency | Cumulative frequency |
| 0 | 3 | 1.1 | 1.1 | 35 | 56.5 | 56.5 |
| 1 | 7 | 2.5 | 3.5 | 5 | 8.1 | 64.5 |
| 2 | 2 | 0.7 | 4.3 | 1 | 1.6 | 66.1 |
| 3 | 13 | 4.6 | 8.9 | 5 | 8.1 | 74.2 |
| 4 | 13 | 4.6 | 13.5 | 8 | 12.9 | 87.1 |
| 5 | 26 | 9.2 | 22.7 | 0 | 0 | 87.1 |
| 6 | 40 | 14.2 | 36.9 | 3 | 4.8 | 91.9 |
| 7 | 50 | 17.7 | 54.6 | 3 | 4.8 | 96.8 |
| 8 | 45 | 16 | 70.6 | 2 | 3.2 | 100 |
| 9 | 38 | 13.5 | 84 | -- | -- | -- |
| 10 | 45 | 16 | 100 | -- | -- | -- |

#### Discriminant validity

The ChaLKS-Medical scores enabled good discrimination between knowledge among healthcare professionals and potential aid workers. The mean number of correct answers among non-medical respondents was 1.8 (standard deviation [SD] 2.4), compared to 7.0 (SD 2.3) among health professionals (T-Test 16.03, p<0.001). The scores for the ChaLKS-Social&Health also discriminated between the two main groups of respondents (non-medical: 1.8 [SD] 2.5 versus 6.9 [SD 1.6], T-test 20.2, p < 0.001).

### Knowledge on Chagas disease

**Table 4** shows the itemized results for the ChaLKS-Medical and ChaLKS-Social&Health questionnaire. The mean number of correct answers on the ChaLKS-Medical among healthcare students and residents was 7.0. Two difficult questions for respondents were: *Can CD be transmitted through blood transfusions?* (67.1% correct answers), and *Can CD be transmitted from mother to child?* (59.0% correct answers). The most frequent mistake among health professionals and medical and pharmacy students related to the name of the drug used to treat CD (23% correct answers).

**Table 4.** Description and results of the questionnaire among the respondents to the ChaLKS-Medical (n = 283) and the ChaLKS-Social&Health (n = 66)

| Items | ChaLKS-Medical<br>(n=283) | ChaLKS-Social&Health (n=66) |
| --- | --- | --- |
| Mean n (SD) correct answers | 7.0 (2.3) | 1.8 (2.4) |
| Is CD endemic to the Americas? n (%) | 257 (90.8) | 13 (19.7) |
| Is CD endemic to Africa? n (%) | 184 (65) |  |
| Is CD endemic to Europe? n (%) | 257 (90.8) | 21 (31.8) |
| Can CD be transmitted through blood transfusions? n (%) | 190 (67.1) |  |
| Can CD be transmitted from mother to child? n (%) | 167 (59.0) | 10 (15.2) |
| Can CD be transmitted through sexual contact? n (%) |  | 13 (19.7) |
| Can CD be transmitted by living with a person who has the disease? n (%) |  | 15 (22.7) |
| Can CD be transmitted by an insect bite? n (%) |  | 27 (40.9) |
| Can CD be transmitted by kissing bugs? n (%) | 192 (67.8) |  |
| Can CD affect the heart? n (%) | 239 (84.8) | 13 (19.7) |
| Can CD affect the stomach and bowels? n (%) | 213 (75.3) |  |
| Is CD diagnosed via serology? n (%) | 214 (75.6) |  |
| Is there treatment for CD? n (%) |  | 22 (33.3) |
| What drug is used to treat CD? n (%) | 66 (23.3) |  |

CD: Chagas disease.

On the ChaLKS-Social&Health scale, potential aid workers responded correctly to an average of 1.8 questions. The question garnering the most mistakes was *Can CD be transmitted from mother to child?* (15.2% correct answers). Respondents also had little knowledge on the endemicity of the disease in the Americas or sexual transmission (19.7% correct answers).

**Table 5** presents the number of ChaLKS-Medical items yielding correct answers among healthcare professionals; those who had received information about CD in the past showed a better average performance (7.2 versus 4.3 correct answers; p < 0.001). This difference was apparent in 9 of the 10 items analyzed (the exception was for the question *Can CD be transmitted from mother to child?*). The same occurred in the ChaLKS-Social&Health questionnaire: those who had received information about CD in the past obtained better scores on the 8-item questionnaire (5.1 versus 1.1 correct answers; p < 0.001). This difference held for all items of the questionnaire.

**Table 5.**
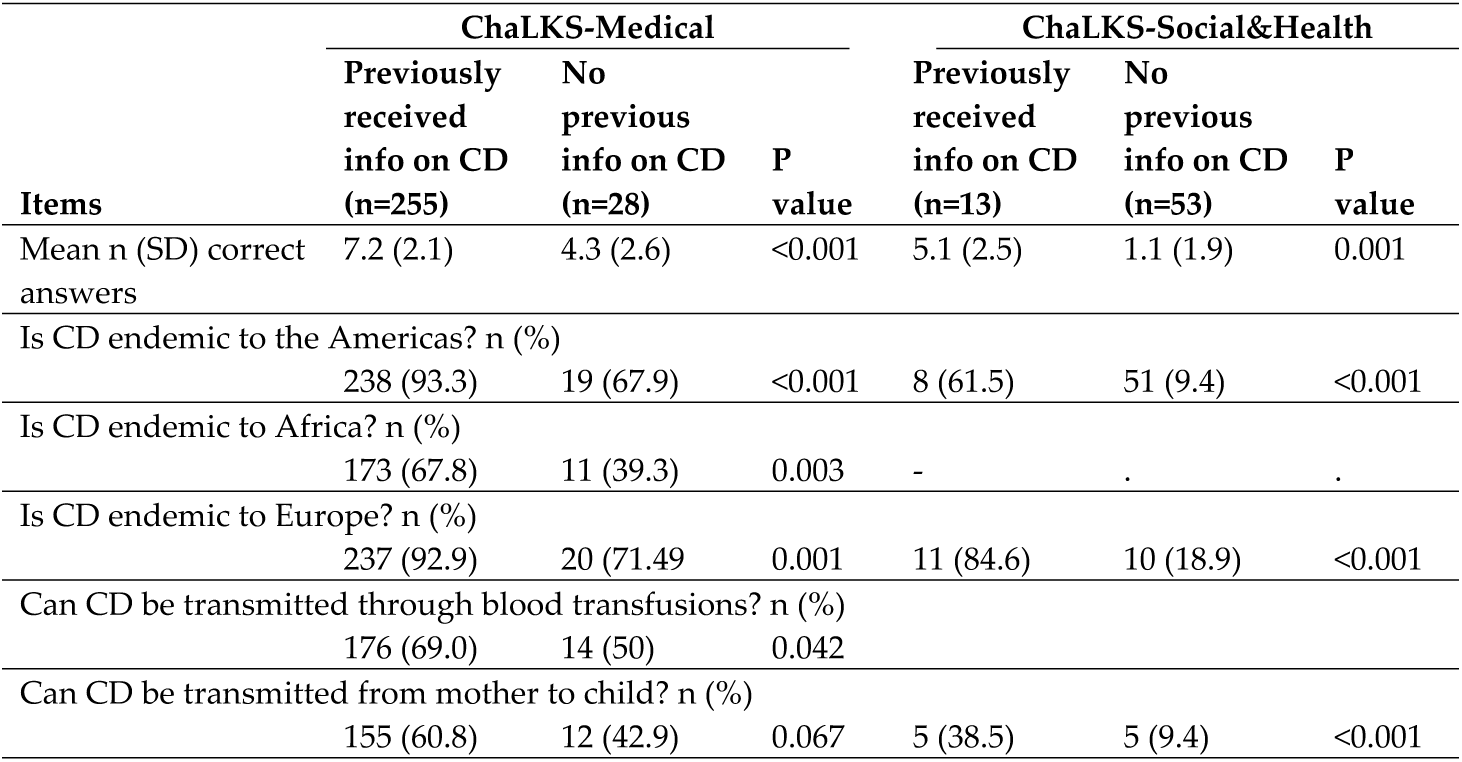
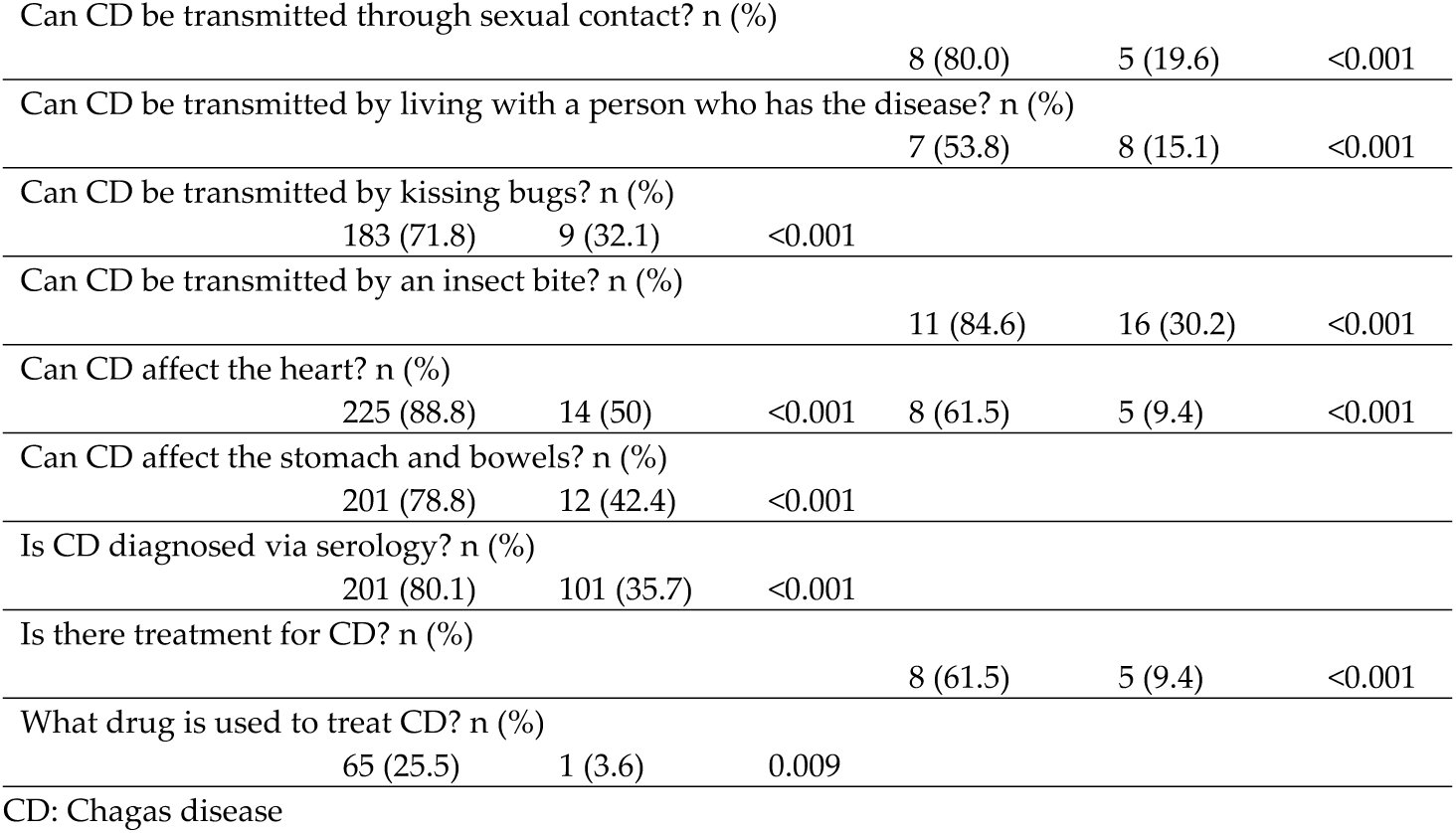
Subanalysis in the respondents to the ChaLKS-Medical (n = 283) and the ChaLKS-Social&Health (n = 66) according to whether they had received previous information on Chagas disease (CD)

The mean number of correct answers among the medical students was 5.8, similarly to the pharmacy students (5.8). In contrast, medical residents scored higher on average (7.4, p < 0.001), and pharmacy residents did comparably well (7.61, p < 0.001). Itemized comparisons among these groups are presented in the supplementary material (**Supplementary material: table S7**).

We also compared responses among the 200 participating medical professionals and pharmacists (ChaLKS-Medical) according to whether they had been born and educated (until their bachelor’s degree) in Latin America versus Europe (including Spain). The number of correct answers was similar in both groups; the only item showing differences was related to the endemicity of CD in Africa, with Latin American professionals performing worse on this item than Europeans (39.2% versus 76.3% correct answers; p < 0.001; **Supplementary material: table S8**).

## DISCUSSION

The statistics applied to the validation analysis of these instruments confirm that both scales have a satisfactory factorial function, internal consistency, adequate difficulty indexes, and capacity to discriminate between different levels of knowledge on Chagas disease. Both instruments are appropriate for supporting aid actions in Chagas-endemic countries or in non-endemic countries providing health and social services to Latin American immigrants.

In general, the level of knowledge shown in response to the questionnaires is adequate, but improvable. Our results were consistent with similar studies performed among physicians and medical students in non-endemic countries [7, 8, 11, 17]. Nevertheless, we drew conclusions based on results in convenience samples from selected populations with a special interest in global health, so our results are not generalizable [13]. Moreover, we speculate that general awareness among physicians of other European countries (where CD is not as prevalent as in Spain) may be even lower [5].

Some key aspects were well known among medical students and healthcare professionals, such as the endemicity of CD in Latin America. Nevertheless, some believed CD was also endemic to Africa, probably because they confused the CD parasite with the phylogenetically related parasite (also belonging to the genera *Trypanosoma*) responsible for African trypanosomiasis. Another epidemiologically crucial aspect was unknown among many healthcare professionals (up to 40%): the mother-to-child transmission route. This is the most important transmission route in non-endemic countries where blood and organ donors are screened. Our results suggest that medical schools and post-graduate medical training should emphasize this aspect to a greater extent, especially in countries with Latin American populations.

Another aspect which respondents struggled with was the name of the drugs indicated for treating CD; less than 30% of the healthcare professionals knew the names, and less than 9% of medical and pharmaceutical students. Nevertheless, we consider it much more important for professionals to know that a treatment exists than to be able to name the drug as such. In our previous pilot study conducted among medical students, 87% knew there was a specific treatment for CD, and around 75% were aware that most of the carriers of *T. cruzi* are asymptomatic [12]. This issue is crucial in order to screen and encourage the population at risk to perform the diagnostic test. If health professionals are aware of the endemicity of the infection, its transmission routes (especially those at play in the country where the professional is based), its asymptomatic nature, which diagnostic test to order if CD is suspected, and the fact that treatment exists, we consider that the health professional is equipped with the basic knowledge to suspect, prevent, and treat CD.

The reactive item, *Can someone with CD feel okay (asymptomatic)?*, which we consider relevant to discriminate the level of knowledge on CD, yielded a very low difficulty index (a high number of correct answers) and a low factorial saturation (0.29) in the analysis of construct validity. In light of these statistics, we chose not to retain the question in the ChaLKS.

As pointed out in other articles, it is very important to measure the level of knowledge of NTDs in undergraduate medical students in order to evaluate the need to intensify education in these concepts, providing future medical doctors with optimal tools to cope with NTDs [21]. We did not observe differences in the level of knowledge among resident physicians trained in Europe versus Latin America. In our opinion, this may reflect an acceptable training in imported diseases in Europe, equaling European medical students’ CD knowledge to the ones performing their bachelor’s degree in endemic areas.

This study has some limitations. One is the low number of students that filled in the ChaLKS-Social&Health. With regard to the items included on the scale, it would have been advisable to add a question on CD transmission during organ transplantation to the initial version of the questionnaire in order to assess how it performed during the validation. Other knowledge items to consider assessing on future questionnaires would be the risk of stroke associated with CD (as some studies show the lack of knowledge on this aspect among health professionals [22]) and the existence of screening programs and activities to control the transmission of *T. cruzi* (regarding pregnant women in endemic areas, blood donations, and organ transplants) in the countries of the surveyed health professionals [1]. This latter aspect would be advisable to include on both versions, given the importance of doctors having up-to-date information on these programs and of other health and social service providers knowing where to refer people who are at risk of carrying *T. cruzi*.

The ChaLKS-Medical includes 10 items with good metric properties: 3 on epidemiology, 3 on transmission, 2 on clinical characteristics, 1 on diagnosis, and 1 on treatment. The question *Is CD endemic to the Americas?* shows high factorial saturation and a low difficulty index, so it could have been excluded from the questionnaire. However, the consulted experts considered this question too relevant to leave out, and it was thus maintained in the final version.

In the validation of the ChaLKS-Medical, pharmaceutical students and professionals were included because their training in parasitic diseases like CD is intense. Moreover, professionals from this sector may have a role in developing diagnostic methods and pharmacological treatments, as well as participating in multidisciplinary hospital teams that treat patients with CD [23].

ChaLKS-Social&Health is an easier scale than ChaLKS-Medical, as its target audience has a different profile. We believe in contains the minimum number of items on CD that any student in social or health studies who is interested in national or international aid work with Latin Americans should know. Our results show that potential aid workers who have previously received information on CD have a higher level of knowledge on the disease, highlighting the desirability of providing enhanced training to this young collective. Approximately 500 aid programs in Latin America are launched from Spain every year [15]. We believe that reinforcing knowledge about CD would be especially useful in the framework of these programs, as it would be beneficial for social and health aid workers aimed at Latin Americans—whether at home or abroad—to have knowledge on CD, regardless of whether the aid projects are directly related to this disease. Not taking this step would be a waste of a good opportunity: knowledge on CD in at-risk populations is low [24]; many people are affected by more than one NTD [25], and integrated programs seem to be effective towards reducing the burden of NTDs in low- and lower-middle-income countries [26].

ChaLKS-Medical also showed a higher rate of correct answers among those who had received previous information on CD, prompting a reflection on the need to continue improving training on CD in healthcare students and professionals, both in endemic and non-endemic countries [12, 13, 27]. Awareness and a precise knowledge of CD in medical students, resident physicians and healthcare professionals in general may facilitate an early diagnosis and correct clinical management in patients with *T. cruzi* infection and greater adherence to *T. cruzi* screening programs [28], also improving the overall healthcare for this population and the control of the disease [29]. Future studies should assess the impact of these data on CD underdiagnosis, evaluating the need for educating students and healthcare professionals from non-endemic countries about this and other emerging diseases [17].

In conclusion, a validated questionnaire on knowledge about Chagas disease can be useful:

1. to use as a teaching tool before and after specific training sessions on CD for doctors, medical students, other health and social professionals, and potential aid workers;
2. to evaluate the level of knowledge among our health professionals and students in areas not endemic to CD, as a step to inform the design of training courses on NTDs and global health;
3. to assess knowledge on CD in aid workers who wish to carry out their work in countries with vector-borne transmission of CD or with Latin American migrants.

It would be desirable that similar initiatives are replicated in other countries and in relation to other NTDs to improve the training in global health among health professionals and aid workers.

## Supplementary material

**Table S1.** Surveyed participants and subgroups

**Table S2**. Preliminary questionnaire with 26 reactive items testing knowledge of Chagas disease (CD)

**Table S3a.** Comparison of the versions of the Spanish language scales for medical/pharmacy students and residents (23 items) and for potential aid workers (15 items)

**Table S3b.** The English-language version for medical/pharmacy students and residents included 23 items and for potential aid workers, 15 items.

**Table S4**. Items included in ChaLKS-Medical and ChaLKS-Social&Health, and key items according to experts

**Table S5a**. Chagas Disease Level of Knowledge Scale for healthcare professionals (ChaLKS-Medical), in Spanish

**Table S5b**. Chagas Disease Level of Knowledge Scale for healthcare professionals (ChaLKS-Medical), in English

**Table S6a**. Chagas Disease Level of Knowledge Scale for potential aid workers of social and healthcare fields (ChaLKS-Social&Health), in Spanish

**Table S6b**. Chagas Disease Level of Knowledge Scale for potential aid workers of social and healthcare fields (ChaLKS-Social&Health), in English

**Table S7.** Subgroup analysis in 283 professionals surveyed on knowledge of Chagas disease, with the scale for healthcare professionals (ChaLKS-Medical), according to occupation

**Table S8**. Subgroup analysis in 200 doctors and pharmacists surveyed on knowledge of Chagas disease, with the scale for healthcare professionals (ChaLKS-Medical, version 2), according to nationality and location of studies in Europe (incl. Spain) or Latin America

## Acknowledgments

We acknowledge Meggan Harris for proofreading of the manuscript, Tania González Zivkovic for the forward-back-translation, and the consulted experts on Chagas disease for their valuable comments.

## Funding

No external funding supports this study.

## Transparency declarations

None to declare.

## Authors’ contribution

Conceptualization: José M. Ramos-Rincón, Miriam Navarro.

Formal analysis: José M. Ramos-Rincón, José J. Mira-Solves, Miriam Navarro.

Investigation: José M. Ramos-Rincón, Violeta Ramos-Sesma, Diego Torrús-Tendero.

Methodology: José M. Ramos-Rincón, José J. Mira-Solves, Jara Llenas-García, Miriam Navarro.

Writing – original draft: José M. Ramos-Rincón, José J. Mira-Solves, Jara Llenas-García, Miriam Navarro.

Writing – review & editing: José M. Ramos-Rincón, José J. Mira-Solves, Violeta Ramos-Sesma, Diego Torrús-Tendero, Jara Llenas-García, Miriam Navarro.

